## Supplemental Figures 1-5 for "Genome-wide CRISPR screens identify novel regulators of wild-type and mutant p53 stability"

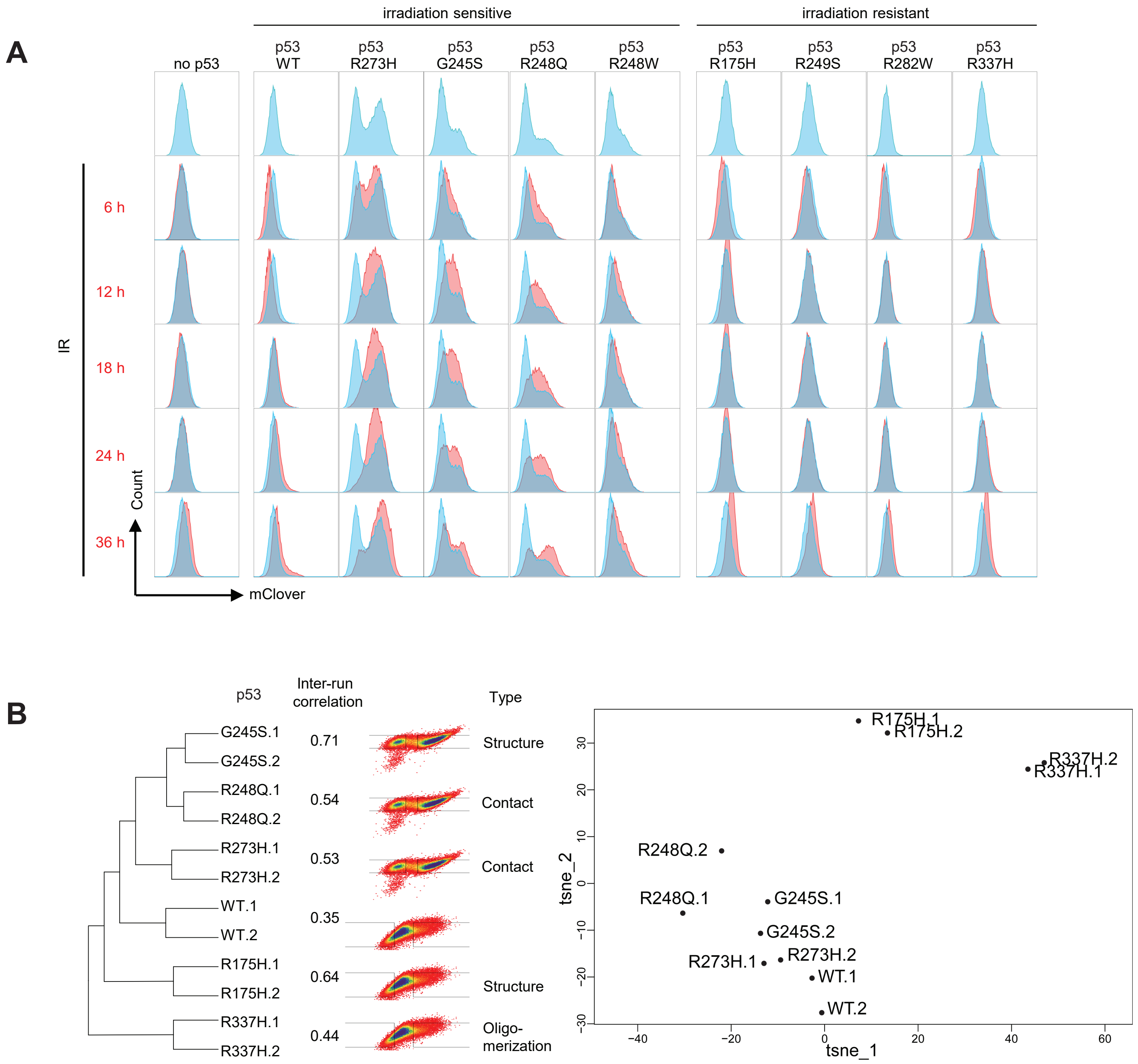


Figure S1. CRISPR screen for regulators of p53 stability.

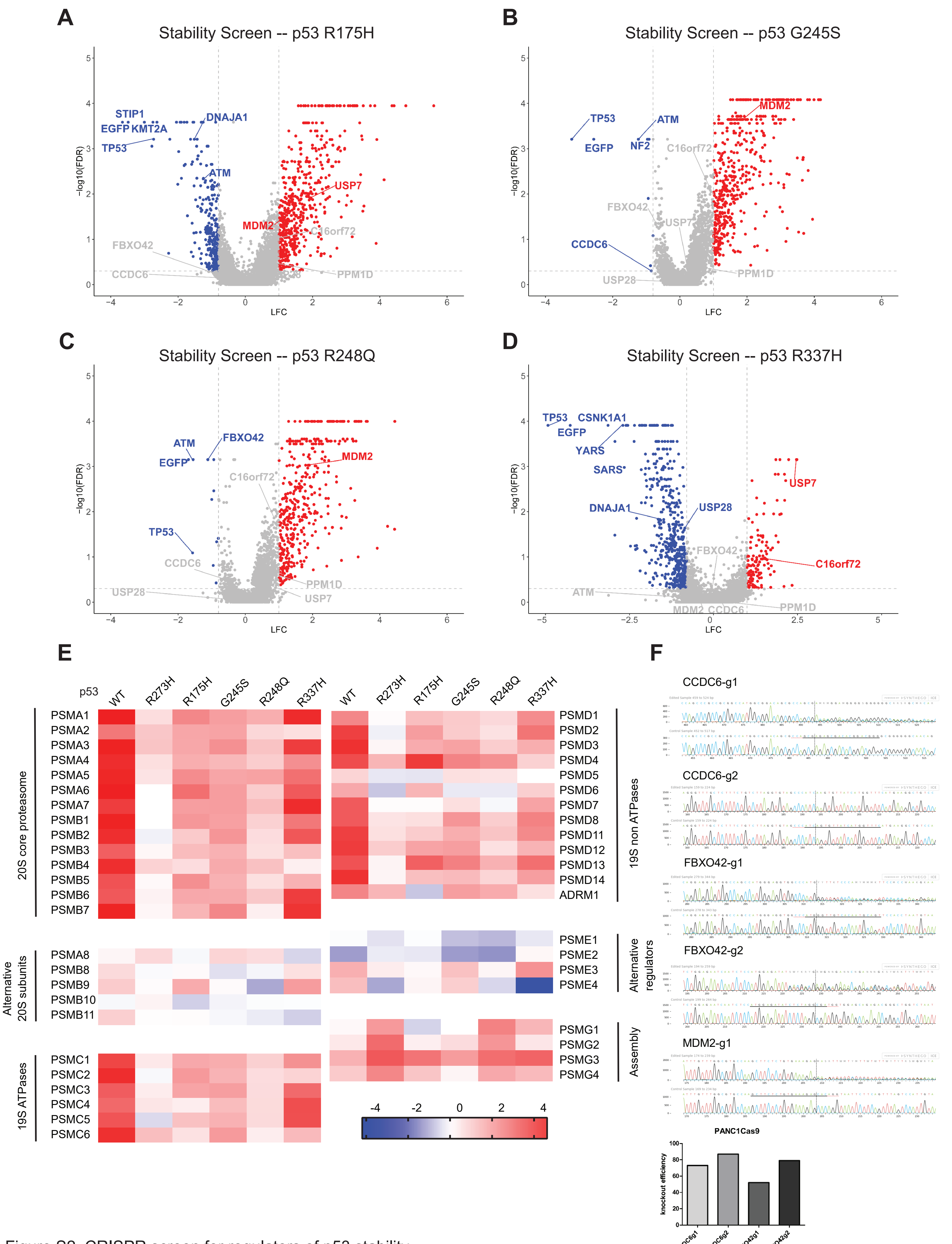


Figure S2. CRISPR screen for regulators of p53 stability.

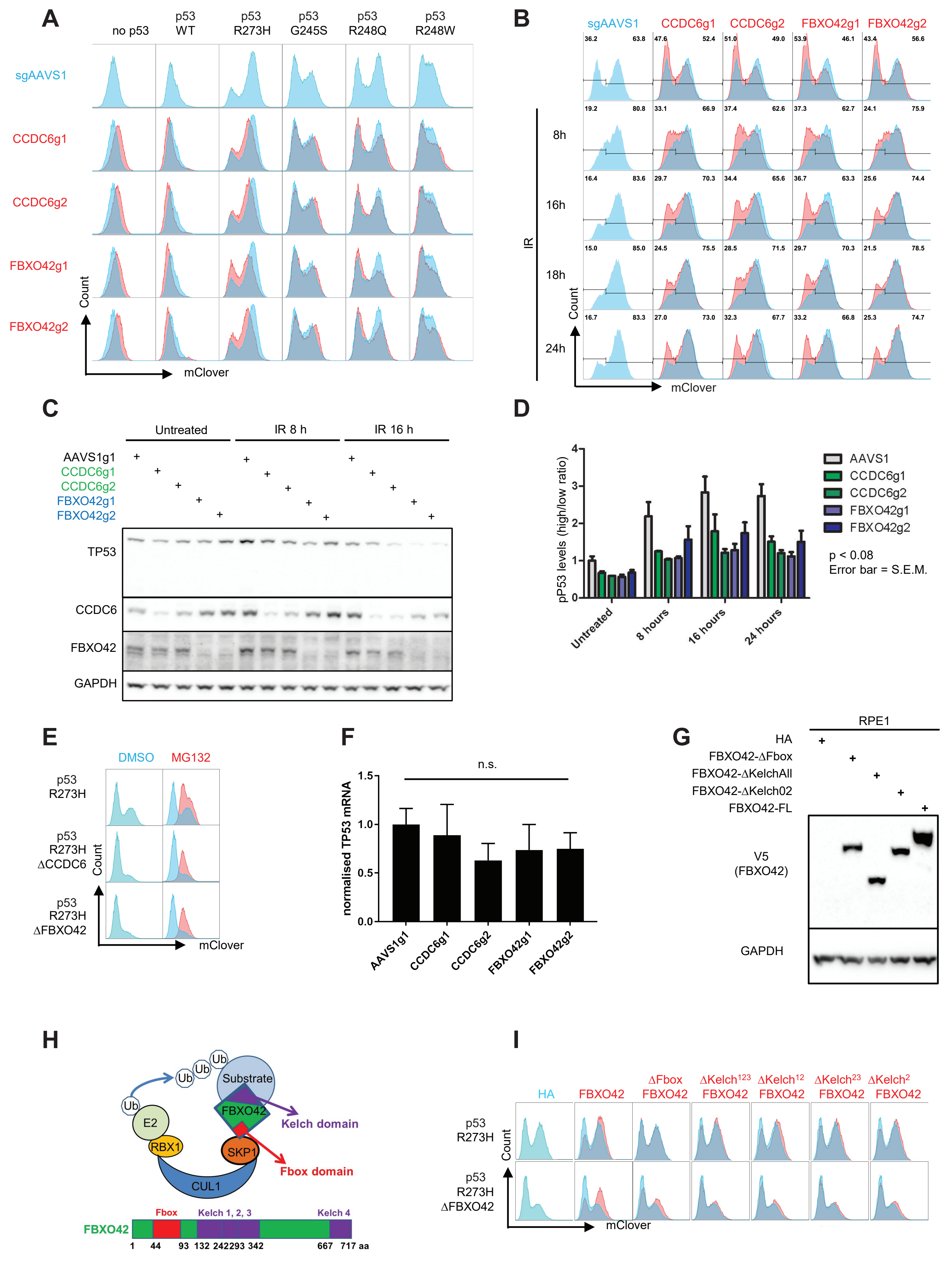


Figure S3. FBXO42-CCDC6 axis regulates p53 stabilities across wild-type and different mutants.

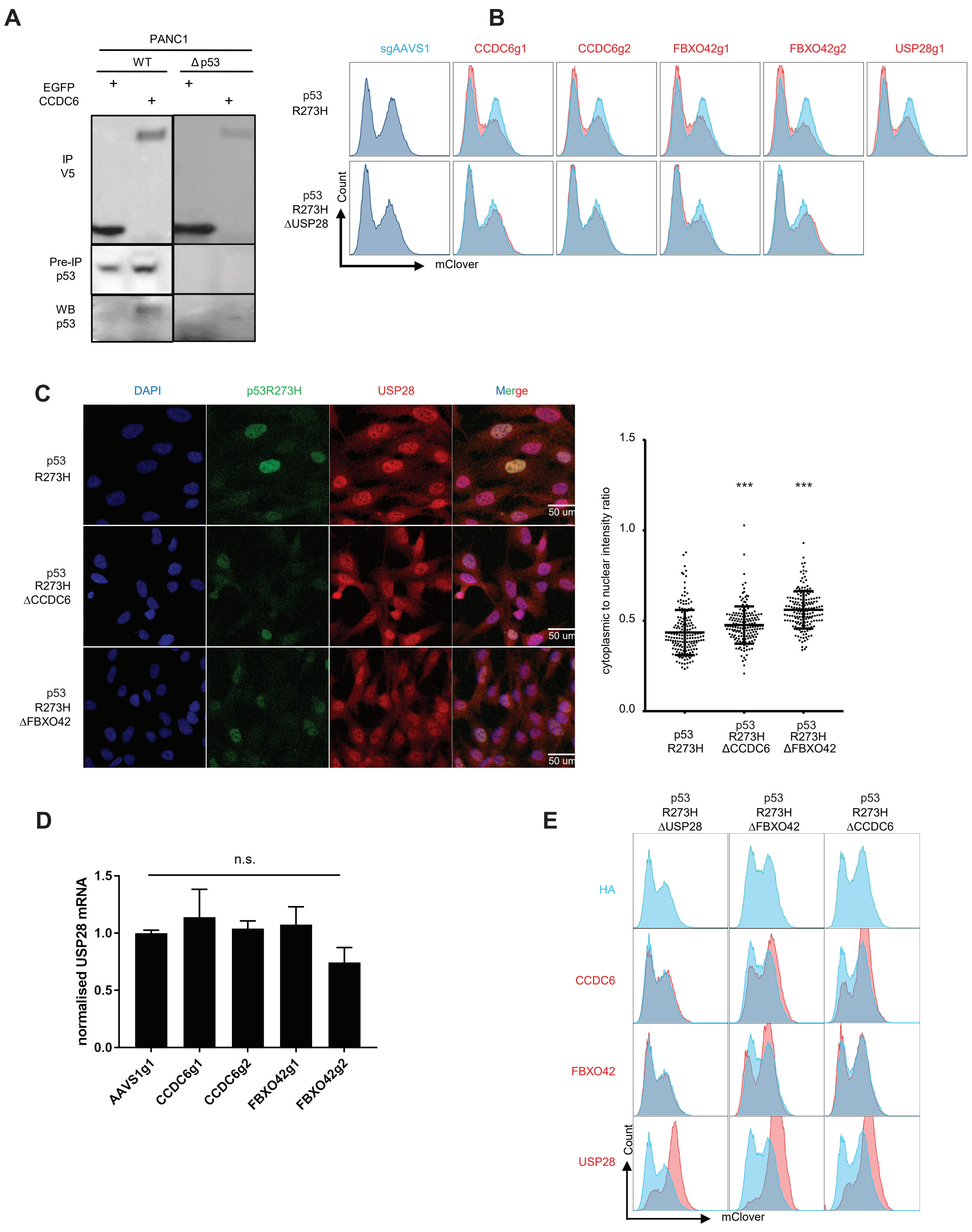


Figure S4. Mapping the genetic interaction network of FBXO42-CCDC6 and mutant p53.

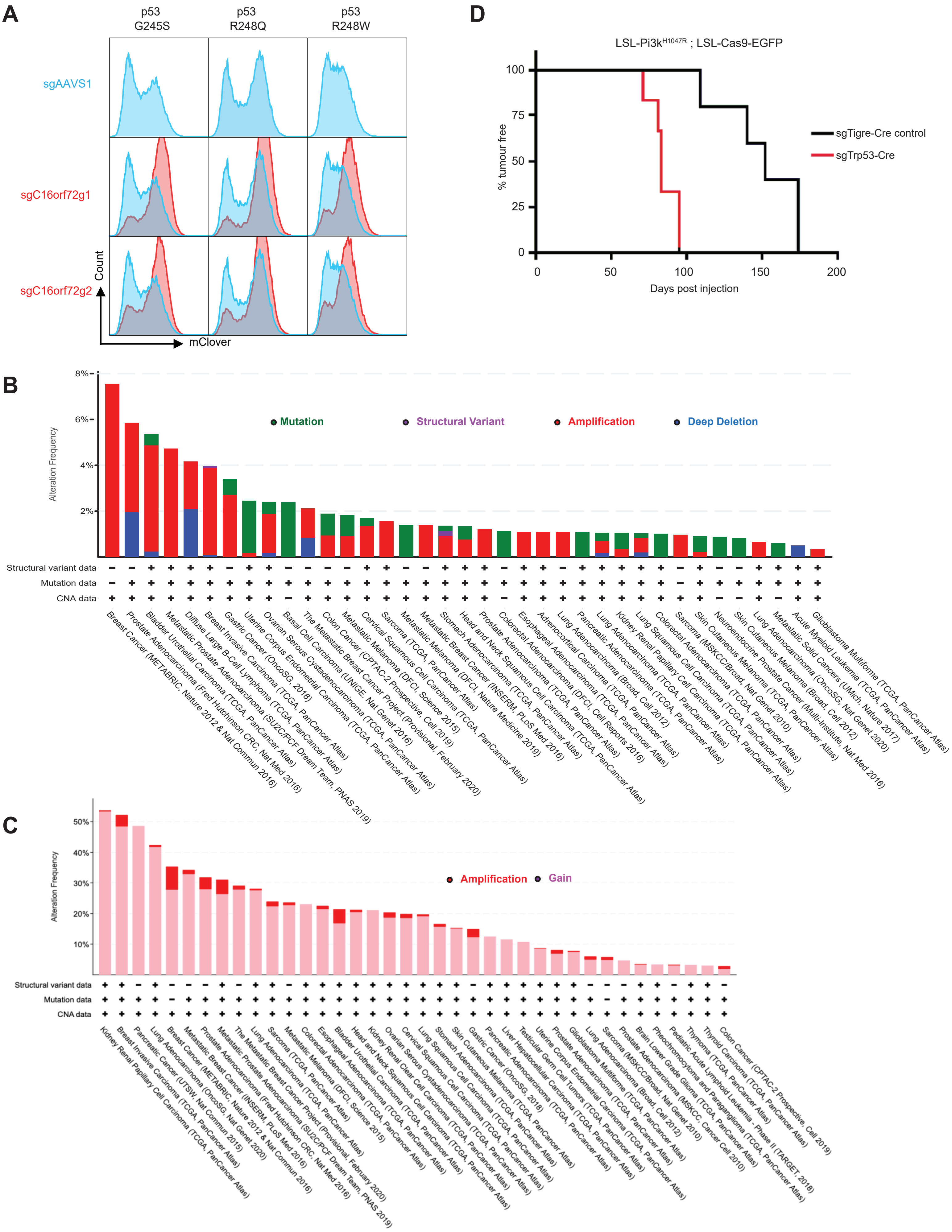


Figure S5. C16orf72 functions as oncogene and regulates p53 stability in the mammary gland.
